# Challenges for error-correction coding in DNA data storage: photolithographic synthesis and DNA decay

**DOI:** 10.1101/2024.07.04.602085

**Authors:** Andreas L. Gimpel, Wendelin J. Stark, Reinhard Heckel, Robert N. Grass

**Author notes:** Correspondence and requests should be addressed to Robert N. Grass.

## Abstract

Efficient error-correction codes are crucial for realizing DNA’s potential as a long-lasting, high-density storage medium for digital data. At the same time, new workflows promising low-cost, resilient DNA data storage are challenging their design and error-correcting capabilities. This study characterizes the errors and biases in two new additions to the state-of-the-art workflow in DNA data storage: photolithographic synthesis and DNA decay. Photolithographic synthesis offers low-cost, scalable oligonucleotide synthesis but suffers from high error rates, necessitating sophisticated error-correction schemes, for example codes introducing within-sequence redundancy combined with clustering and alignment techniques for retrieval. On the other hand, the decoding of oligo fragments after DNA decay promises unprecedented storage densities, but complicates data recovery by requiring the reassembly of full-length sequences or the use of partial sequences for decoding. Our analysis provides a detailed account of the error patterns and biases present in photolithographic synthesis and DNA decay, and identifies considerable bias stemming from sequencing workflows. We implement our findings into a digital twin of the two workflows, offering a tool for developing error-correction codes and providing benchmarks for the evaluation of codec performance.

## Introduction

Many error-correction codes have been developed and optimized for DNA data storage in an effort to showcase DNA’s potential as a long-lasting, high-density storage medium for digital data.^1–6^ Since its first successful demonstrations in vitro by Church et al.^7^ and Goldman et al. ^4^, significant improvements in the fidelity of DNA synthesis and the construction of error correction codes have enabled DNA data storage to come close to its theoretical limits in terms of code rate and storage density.^1,8^ In doing so however, many error-correction codes are designed for low-error, low-bias scenarios involving high-fidelity DNA synthesis and sequencing, without aging-induced DNA decay.^1,9,10^ As a result, two important new additions to this state-of-the-art DNA data storage workflow currently lack established and optimized implementations of error-correction codes: photolithographic synthesis and DNA decay (see Fig. 1).

**Fig. 1:**
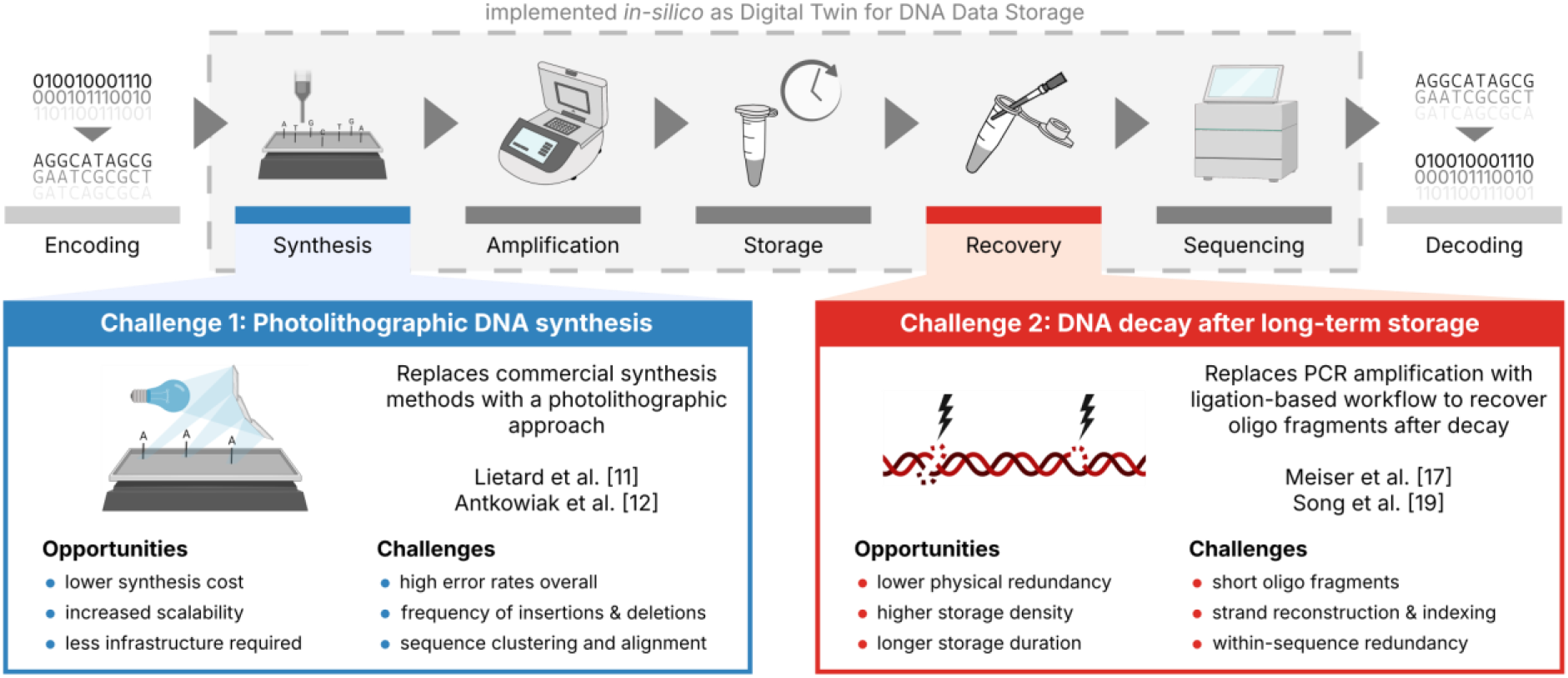
Current challenges in the DNA data storage workflow. Illustration of the current state-of-the-art DNA data storage workflow (top) with the two new additions discussed in this study highlighted: photolithographic synthesis (blue box) and DNA decay (red box). Both the state-of-the-art workflow and the challenges have been implemented in-silico as a Digital Twin for DNA Data Storage (grey box, see also Gimpel et al.^9^) to support codec development and experimental planning.

As an alternative to commercial DNA synthesis, photolithographic DNA synthesis promises low-cost and scalable synthesis of oligonucleotide pools at the cost of high error rates,^8,11,12^ as demonstrated by Lietard et al.^11^ and Antkowiak et al.^12^ (see Fig. 1, blue box). An important application of photolithographic synthesis lies in enabling DNA-of-things architectures,^13,14^ as high storage densities and low physical redundancies are not needed in this application. Instead, low synthesis costs are critical to the architecture’s adoption as a tool to embed information into objects.^13–15^ For this reason, photolithographic synthesis is an interesting challenge for error-correction coding, as the high physical redundancy (e.g., >200x) and sequencing depth (e.g., >50x) can be exploited to generate less erroneous consensus sequences by effective clustering, multiple-sequence alignment, and majority voting.

Recovery of oligos after long-term storage is commonly performed via PCR amplification using adapters included in the sequences.^16^ As this recovers only intact, full-length oligos – a small fraction of the available sequence information^17^ – DNA decay is commonly limiting the effective storage density (i.e., code rate / physical coverage) by requiring high physical coverage.^9,10,18^ Thus, achieving storage densities close to the theoretical limit^7^ of 455 EB g^-1^ will require data recovery from low physical coverages rather than marginally improved code rates. To this end, the decoding of oligo fragments after DNA decay is a viable option, at the cost of increased complexity during decoding,^10,17,19^ as shown by Song et al.^19^ and Meiser et al.^17^ (see Fig. 1, red box). Specifically, the capability of data recovery from oligo fragments – either by their assembly into full-length sequences or their partial decoding – presents an opportunity for error-correction codes to realize the high storage density afforded by low physical coverage (demonstrated as low as 10x by Organick et al.^20^).

In this study, we characterize the errors and biases present in two new additions to the state-of-the-art workflow in DNA data storage – photolithographic synthesis and DNA decay (see Fig. 1) – and highlight the associated challenges for error-correction codes. To do so, we use analytical tools previously developed for DNA data storage^9^ with existing sequencing data from literature.^11,12,17,19^ In discussing the observed error patterns and biases, we identify opportunities for the optimization of experimental workflows and the development of error-correction codes. Finally, we define two challenges – realistic scenarios for current challenges in DNA data storage involving photolithographic synthesis or DNA decay – and provide the corresponding tools for testing and benchmarking implementations of error-correction codes for these challenges.

## Results

### Photolithographic synthesis yields highly erroneous DNA

For the first challenge, we performed a detailed re-analysis of the error patterns produced during photolithographic DNA synthesis, based on the sequencing data by Lietard et al.^11^ and Antkowiak et al.^12^ (four and three datasets respectively, see Methods). As shown in Fig. 2a, the mean error rate in the sequencing data exceeded 0.1 errors per nucleotide (nt^-1^) for all datasets, with up to around 0.2 nt^-1^ for the high-density syntheses by Antkowiak et al.^12^ (datasets *File 2* and *File 3*). Therefore, in the best case, about one in every 10 nucleotides was erroneous after photolithographic DNA synthesis. These error rates are in-line with the analysis by the original authors,^11,12^ and in stark contrast to the error rates commonly observed for established commercial DNA synthesis, which usually lie below 0.02 nt^-1^ even in the worst case.^9,18^ Interestingly, deletion errors dominate both in photolithographic and commercial DNA synthesis.^9,18,21^ However, similarly to electrochemical synthesis in which a failure to deprotect the growing oligo causes a deletion,^22,23^ part of the high error rates during photolithographic synthesis could be attributed to insufficient deprotection during the illumination step. The two aforementioned high-density syntheses by Antkowiak et al.^12^ did not show the same error distribution, as substitution errors were dominating instead (see *File 2* and *File 3* in Fig. 2a).

**Fig. 2:**
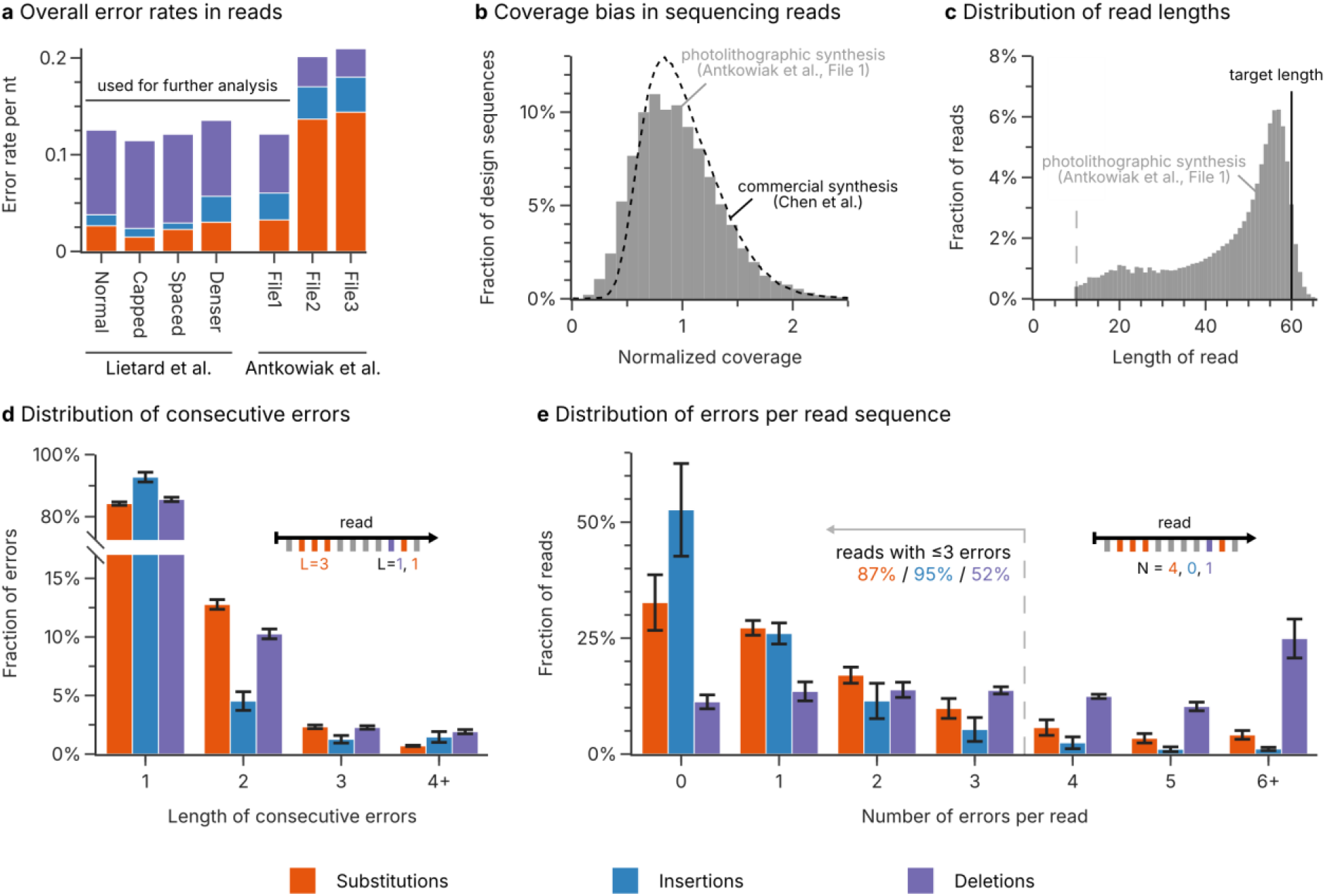
Analysis of photolithographic synthesis. **(a)** Overall rate of substitution (orange), insertion (blue), and deletion (purple) errors in the sequencing datasets for photolithographic syntheses by Lietard et al.^11^ and Antkowiak et al.^12^. The error rates represent the median error rate across the length of the sequence, to minimize the effect of the low-diversity regions at the start and end of the sequences. **(b)** Distribution of the sequence coverage after photolithographic synthesis (Antkowiak et al.^12^, File 1, grey bars) compared to the coverage distribution for a commercial synthesis by material deposition (Twist Biosciences, data by Chen et al.^24^). **(c)** Length distribution of the reads after photolithographic synthesis (Antkowiak et al.^12^, File 1, grey bars) compared to the length of the design sequences (solid line). Only the segment of the read which aligned to the reference sequence is considered. Reads smaller than 10 nucleotides were discarded during mapping of the sequencing data (dotted line). **(d)** The length of consecutive substitution (orange), insertion (blue), and deletion (purple) errors in the sequencing datasets for photolithographic syntheses by Lietard et al.^11^ and Antkowiak et al.^12^ (only File 1). **(e)** The number of substitution (orange), insertion (blue), and deletion (purple) errors in the sequencing reads for photolithographic syntheses by Lietard et al.^11^ and Antkowiak et al.^12^ (only File 1). The bars in **(d+e)** show the mean fraction across the four datasets, with the standard error of the mean shown as error bars. The insets illustrate how the length of consecutive errors and the number of errors per read are evaluated.

Given the two error regimes present in the datasets on photolithographic synthesis, we chose to use the low-error datasets (i.e., all four datasets of Lietard et al.^11^ and *File 1* by Antkowiak et al.^12^) as a best-case scenario for further analysis. These datasets featured error rates of 0.082 deletions, 0.016 insertions, and 0.025 substitutions per nucleotide on average, representing the current state-of-the-art for photolithographic DNA synthesis. Assessing the occurrence of consecutive errors (see Fig. 2d) revealed that error events spanning multiple consecutive nucleotides were more common than expected. However, they still only represented a minority: about 16% of substitutions, 14% of deletions, but only 7% of insertions occur in groups of at least two consecutive errors. In contrast, the number of independent errors per read followed the expected binomial distribution (see Fig. 2e). Due to the high error rates of photolithographic synthesis, only around 2% of sequencing reads were fully error-free (compared to up to 81% of reads for commercial synthesis)^9^ and only 52% of reads contained fewer than four deletions (see Fig. 2e). Considering the short length (60-76 nt) of the synthesized sequences, this high frequency of deletion errors is also a major contributor to the broad distribution of read lengths observed in the data, as shown in Fig. 2c.

### Photolithographic synthesis shows low coverage bias

Previous studies investigating the homogeneity of oligo pools – whether the design sequences are equally represented among the sequencing reads – have reported moderate to strong biases in some commercial syntheses.^9,13,24,25^ As these biases in sequence coverage can lead to sequence dropout which is costly to correct with redundancy, pool homogeneity must also be evaluated for photolithographic synthesis. As shown in Fig. 2b with the comparison of one photolithographic synthesis of Antkowiak et al.^12^ to the commercial synthesis analysed by Chen et al.^24^, the photographically synthesized pool fortunately exhibited a low bias in sequence coverage, comparable to that after commercial synthesis by material deposition.^9,24^ However, increasing the number of sequences synthesized in parallel appears to lead to a marked decline in pool homogeneity (see Supplementary Fig. 1 for all datasets). Nonetheless, sequence dropout due to strong coverage bias is likely irrelevant for photolithographic synthesis, especially given the high physical redundancy and sequencing depth commonly used for it. Therefore, as described above, the main challenge in error-correction coding for photolithographic synthesis lies in effectively utilizing the redundancy afforded by the high physical coverage and sequencing depth to decrease the extreme error rates to feasible levels.

### DNA decay causes few errors but many short fragments

For the second challenge, DNA decay during long-term storage, we analysed the errors and fragmentation patterns during aging of DNA oligos, based on the sequencing data by Meiser et al.^17^ and Song et al.^19^ (three and four datasets respectively, see Methods). Importantly, the datasets differed both by their storage durations (four weeks at 30°C for Meiser et al.^17^, and 28, 56, and 70 days of aging at 70°C for Song et al.^19^) and their workflow for oligo recovery (Swift Biosciences Accel-NGS 1S Plus^26^ for Meiser et al.^17^, and Illumina TruSeq DNA PCR-Free^27^ for Song et al.^19^). In addition, Meiser et al.^17^ tested the effect of enzymatic repair on their aged sample in a separate sequencing experiment (dataset *repaired*). Despite the differences in storage duration and oligo recovery, all datasets exhibited low overall error rates on the order of 0.01 nt^-1^ and were dominated by substitutions (see Fig. 3a). Confirming our previous analysis,^9^ aging only caused a negligible amount of additional substitution errors and did not affect the homogeneity of sequence coverage at all (see Supplementary Fig. 2).

**Fig. 3:**
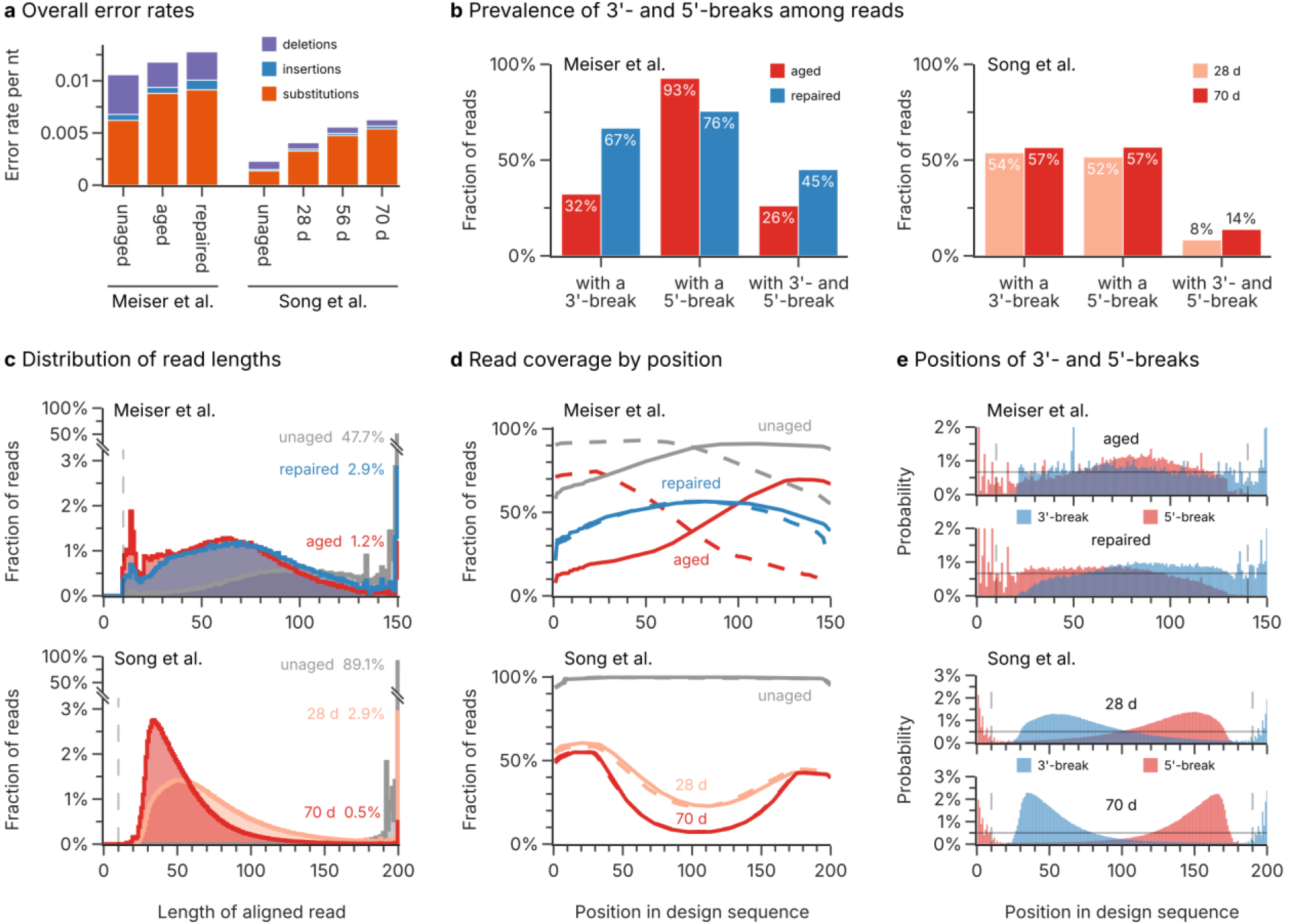
Analysis of DNA decay during aging. **(a)** Overall rate of substitution (orange), insertion (blue), and deletion (purple) errors in the sequencing datasets by Meiser et al.^17^ and Song et al.^19^. **(b)** Fraction of reads with different break patterns in selected datasets by Meiser et al.^17^ (left; red: aged, blue: repaired) and Song et al.^19^ (right; light red: 28 d, dark red: 70 d). **(c)** Histograms of the length distribution of aligned sequencing reads in Meiser et al.^17^ (top; gray: unaged, red: aged, blue: repaired) and Song et al.^19^ (bottom; gray: unaged, light red: 28 d, dark red: 70 d). The fraction of reads with full length in each dataset are given as a percentage. Note that the vertical axis is broken to show the full height of the full-length reads in the unaged datasets. **(d)** Read coverage by position in the design sequence of Meiser et al.^17^ (top; gray: unaged, red: aged, blue: repaired) and Song et al.^19^ (bottom; gray: unaged, light red: 28 d, dark red: 70 d). The reads are separated by their direction into sense (solid lines) and antisense (dashed lines) directions. A value of 100% corresponds to every read containing a specific position. **(e)** Positional distributions of 3’- (blue) and 5’-breaks (red) in selected datasets by Meiser et al.^17^ (top; upper plot: aged, lower plot: repaired) and Song et al.^19^ (bottom; upper plot: 28 d, lower plot: 70 d). The horizontal solid line denotes the expected breakage probability if decay occurs uniformly.

In contrast to photolithographic synthesis (see above), the main challenge for error-correction codes after DNA decay lies in the utilization of oligo fragments, rather than the prevalence of base errors. In addition, the use of ligation-based workflows after DNA decay necessarily leads to an equal proportion of sequencing reads in the sense and antisense directions of the design sequence, further complicating the decoding step. As shown in Fig. 3c, the fraction of full-length sequencing reads drops from 47.7%/89.1% to as low as 1.2%/0.5% for the studies by Meiser et al.^17^ and Song et al.^19^ respectively. Instead, the majority of sequencing reads are around 30-100 nt long, indicating multiple strand cleavages had occurred on each oligo. Notably, all datasets start to show a sharp decline for reads shorter than around 40 nt (see Fig. 3c). This discrepancy can be explained by the use of magnetic beads for clean-up steps in the oligo recovery workflows,^26,27^ which removes short oligos^28^ (see Supplementary Fig. 3).

### Observed breakage patterns after DNA decay show positional bias

Decay is generally assumed to occur uniformly along the length of an oligo,^10,29,30^ leading to two oligo fragments via an hydrolysis pathway^17^ (see Fig. 4a). As a result, breaks at the 3’- or 5’-end of sequencing reads should occur at equal rates, and their positions in the oligos should be uniform. Surprisingly, the observed frequency of 3’- and 5’-breaks in the sequencing datasets deviates significantly from this assumption. As shown in Fig. 4b, the sequencing reads in Meiser et al.’s^17^ aged sample almost universally featured 5’-breaks (93% of reads), but only 32% of reads possessed a break at their 3’-end. While the aged samples of Song et al.^19^ showed no such bias for 5’-breaks, the frequency of double breaks (i.e., both 3’- and 5’-breaks) in the sequencing data of both studies was much lower than expected given the extent of decay (see Fig. 3b).

**Fig. 4:**
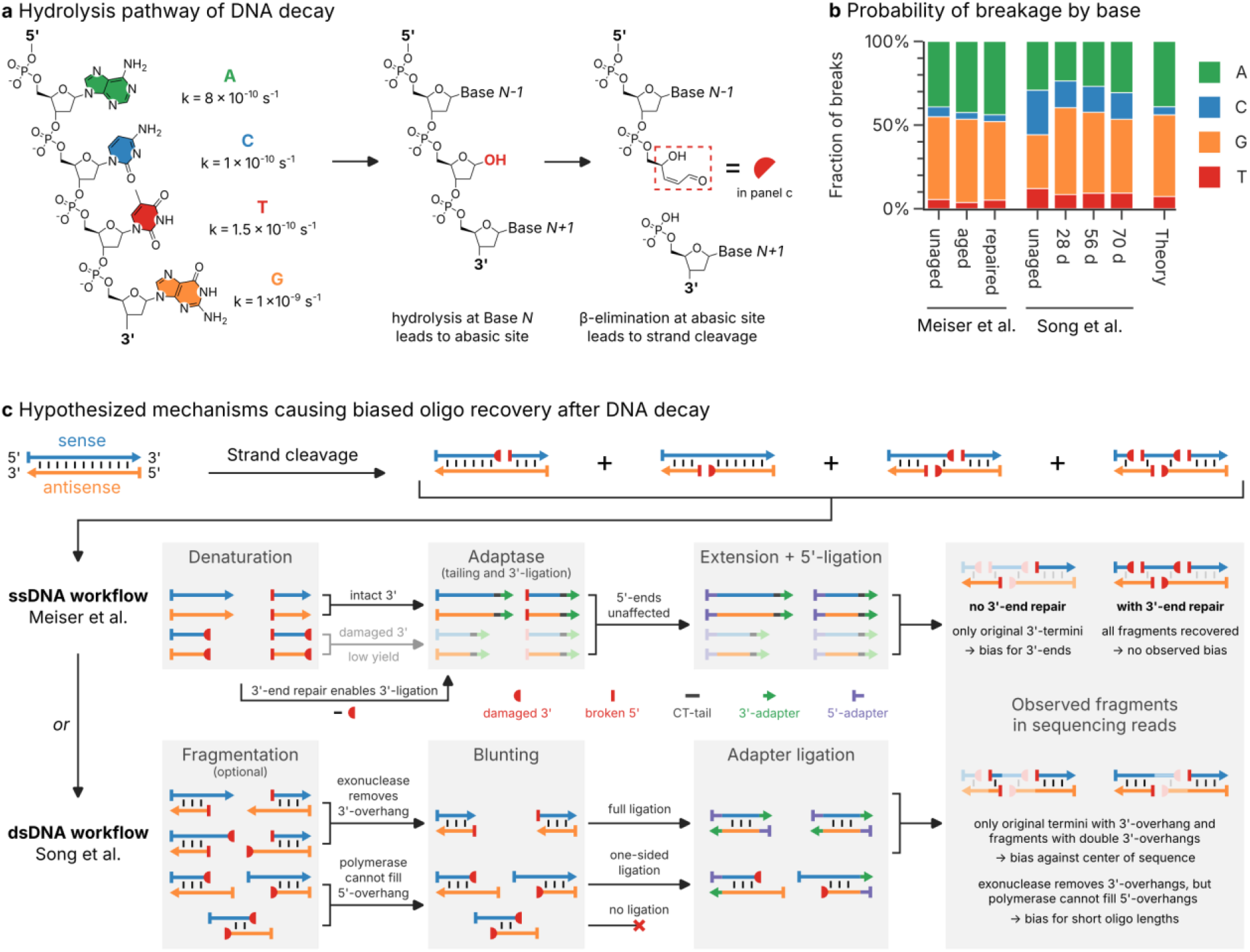
Mechanistic insights into the DNA decay process and the oligo recovery workflow. **(a)** Schematic illustration of the hydrolysis pathway of DNA decay, involving the generation of an abasic site by hydrolysis of the N-glycosidic linkage and subsequent β-elimination to yield two cleaved oligo fragments.^17,35^ Also shown are the approximate rate constants of hydrolysis for each base, at pH 7.4 and 37°C.^37^ **(b)** Distribution of nucleobases (green: A, blue: C, orange: G, red: T) involved in strand cleavage in the sequencing datasets by Meiser et al.^17^ and Song et al.^19^, as well as the theoretical distribution expected from reaction rates (see panel a). Experimental distributions are estimated from the type of nucleobase preceding a 5’-break in the sequencing data. **(c)** Hypothesized mechanisms for the oligo recovery workflows of Meiser et al.^17^ (ssDNA workflow, top) and Song et al.^19^ (dsDNA workflow, bottom). Strand cleavage of sense (blue) and antisense (orange) oligos leads to blocked 3’-ends (3’-PUA, red half-circle) and new 5’-ends (red bar). The ligation of the 3’-adapter (green arrow) and 5’-adapter (violet bar) necessary for sequencing proceeds by different steps in both workflows, leading to unique biases in the sequencing data.

The strong bias in observed fragments is also evident from the read coverage by position and the observed position of 3’- and 5’-breaks, as shown in Fig. 3d+e. The read coverage in the aged dataset from Meiser et al.^17^ is largest at the 3’-ends of sequences, then drops sharply towards the 5’-end (see Fig. 3d, top). Accordingly, the positions of 5’-breaks are biased towards the 3’-end (see Fig. 3e, top). The enzymatic treatment then removed these biases: both the read coverage (see Fig. 3d, top, labelled *repaired*) and the positions of 3’/5’-breaks (see Fig. 3e, top, labelled *repaired*) were close to uniform, with only slight bias against either end. In contrast, the aged datasets of Song et al.^19^ featured a read coverage that was strongly biased against the centre of the sequences (see Fig. 3d, bottom). This is especially evident from the positions of breaks, which were heavily biased towards the 5’- and 3’-end respectively, with the bias worsening with increased storage duration (see Fig. 3e, bottom).

Taken together, the observed fragmentation patterns in the sequencing data do not support the assumption of uniform, independent strand cleavage during DNA decay. However, the differences in fragmentation patterns between the datasets by Meiser et al.^17^ and Song et al.^19^, which differ mostly in the workflow used for oligo recovery (see Methods), suggest bias in the oligo recovery workflow is responsible for these observations. This is also supported by the dramatic reversal to unbiased, uniform breakage patterns after enzymatic treatment in the Meiser et al.^17^ data, indicating a chemical reason in the structure of oligo fragments after decay.

### Strand cleavage is biased towards G and A and yields blocked 3’-ends

In order to understand the mechanism by which the choice of workflow may affect the observed breakage patterns in the sequencing data, we compared the hydrolysis pathway of DNA decay (see Fig. 4a) with the enzymatic steps involved in the workflows by Meiser et al.^17^ and Song et al.^19^ (see Fig. 4c). In the most relevant form of DNA decay, after an abasic site is created via hydrolysis of a glycosidic linkage, the phosphate backbone at this abasic site is rapidly cleaved via β-elimination (see Fig. 4a).^17^ Due to their lower stability against hydrolysis (see rate constants shown in Fig. 4a),^17^ strand cleavage is expected to involve mainly G (49%) and A (39%) rather than T (7.3%) or C (4.9%). This base preference of strand cleavage was accurately reproduced in the sequencing data, with the distribution of nucleotides preceding a 5’-break heavily biased towards G and A in aged samples (see Fig. 4b). This finding supports hydrolysis-induced strand cleavage as the predominant decay mechanism for both storage conditions (30°C and 70°C), but cannot explain the observed positional bias in the breakage patterns.

Besides its nucleobase bias, the hydrolysis pathway of DNA decay yields two new fragments containing the nucleotides up to, and after the abasic site (i.e., without the former nucleobase at the abasic site), each with an intact, phosphorylated 5’-end (see Fig. 4a). However, the new 3’-end created for the formally 5’-part of the oligo is blocked by a 3’-phospho-α, β-unsaturated aldehyde (3’-PUA) remaining after β-elimination of the abasic deoxyribose moiety (red box in Fig. 4a).^17,31,32^ 3’-PUA is known to prevent extension by polymerases in the cellular base excision repair pathway, requiring prior removal of this residue by a phosphodiesterase, for example the apurinic/apyrimidinic endonuclease APE1.^31–35^ The presence of 3’-PUA at the 5’-fragment after cleavage is therefore a possible cause for the observed lack of 3’-breaks in the sequencing data by Meiser et al.^17^, as well as the observed lack of double breaks in the sequencing data of Song et al.^19^.

### Workflow choice and 3’-end repair are crucial for optimal oligo recovery

Further investigation into the effect of the blocking 3’-PUA residue on the individual steps of the oligo recovery workflows helps to explain the observed breakage patterns in Fig. 3b-e fully. As shown in Fig. 4c (ssDNA workflow, top), the workflow used by Meiser et al.^17^ (based on Swift Biosciences Accel-NGS 1S Plus) operates on denatured, single-stranded oligos, and involves tail-mediated 3’-ligation and single-base overhang 5’-ligation of sequencing adapters.^12,26^ As the presence of 3’-PUA in oligo fragments prevents tailing,^35,36^ the 3’-ligation of the sequencing adapter is hindered. Thus, only the fragments with the original, unblocked 3’-ends are recovered. After sequencing, this yields reads with a bias towards the respective 3’-ends of the sense and antisense directions of the design sequence, precisely as was found in the sequencing data by Meiser et al.^17^ (see Fig. 3d+e, top). In their enzymatic repair mix, Meiser et al.^17^ included the enzyme APE1, whose 3′-phosphodiesterase activity repairs the 3’-end by cleaving the 3’-PUA residue.^32–34^ As a result, the tail-mediated 3’-ligation proceeds unhindered and the sequencing data shows a uniform breakage pattern with no positional bias (see Fig. 3d+e, top, labelled *repaired*).

In contrast, the double-stranded oligo recovery workflow of Song et al.^19^ (based on Illumina TruSeq DNA PCR-Free) uses simultaneous blunt-end ligation at the 3’- and 5’-ends after a blunting step with an exonuclease and polymerase.^27^ During blunting, double-stranded oligo fragments with 3’-overhangs – irrespective of whether the 3’-end is intact or blocked – are shortened by the exonuclease activity and are ligated as intended. For fragments with 5’-overhangs however, the polymerase-mediated extension of the recessed 3’-end cannot proceed, due to the 3’-PUA residue.^32,33^ The resulting lack of a blunt end then precludes adapter ligation in the second step. Specifically, only those fragments with the original, blunt-end, as well as fragments with 3’-overhangs at both termini will be recovered. As a result, a strong bias towards short reads close to the design sequences’ 3’- and 5’-ends is expected, precisely as found in the sequencing data by Song et al.^19^ (see Fig. 3c-e, bottom).

These result show that the choice of sequencing preparation for fragmented oligos will introduce considerable bias into the sequencing data. However, the enzyme-treated data from Meiser et al.^17^ presents a viable method to circumvent this bias by adding an enzymatic repair step. Crucially, this means common assumptions for DNA decay (e.g., uniform breakage probability, homogeneous base coverage) are valid and error-correction codes can rely on them.

### Error-correction codes must adapt to challenges

To test the current state-of-the-art of error-correction coding for photolithographic synthesis and DNA decays, we benchmarked three representative codecs from the literature using a digital model of both scenarios implemented in our Digital Twin for DNA Data Storage^9^ (DT4DDS, see Methods and Supplementary Fig. 6+7). For their experiments on photolithographic synthesis, Antkowiak et al. proposed an implementation of the DNA-RS codec developed by R. Heckel^16,38^ with a clustering step.^12,39^ For their experiments on DNA decay, Song et al. used DBGPS, a combination of a fountain code with a de novo strand assembly step in order to reconstruct full-length sequences from fragments.^19,40,40^ In addition, we tested DNA-Fountain by Erlich et al.^3^, the first and most established implementation of fountain codes for DNA data storage. For all codecs, we encoded a 51.6 kB file (Supplementary Fig. 5) using their default (hyper-)parameters, changing only the number and length of generated sequences (see Methods). As benchmarks, we selected the two challenges presented above: first, a use case for photolithographic synthesis in a DNA-of-things architecture;^13,14^ and second, a scenario of DNA decay in long-term archival data storage at the limit of physical redundancy.^20^

The performance of the three error-correction codes in our challenges is shown in Fig. 5. As defined for the challenge on photolithographic synthesis (see Fig. 5, top right, and Methods), high storage density and low physical redundancy are not the major concerns. For this reason, an efficient implementation of error-correction coding for photolithographic synthesis can exploit the high physical redundancy (200x) and sequencing depth (50x) in this scenario to generate less erroneous consensus sequences, for example by clustering, multiple-sequence alignment, and majority voting. Accordingly, the implementation used in Antkowiak et al.^12^ – employing a clustering step and sufficient within-sequence redundancy – outperforms the other codecs, achieving a code rate of 0.6 bit nt^-1^ in Challenge 1. In contrast, our challenge on DNA decay (see Fig. 5, bottom right, and Methods) emphasizes the necessity to utilize oligo fragments for data recovery after long-term storage. Here, the implementation used in Song et al.^19^ – which includes a strand assembly step – achieved a code rate of 1.1 bit nt^-1^, equivalent to an impressive effective storage density of 125 EB g^-1^ (neglecting adapters). In contrast, DNA-Fountain by Erlich et al.^3^ was unable to recover any data in both challenges, despite using code rates as low as 0.2 bit nt^-1^. This is not unexpected, given it was designed for high-fidelity, low bias workflows, like many other error-correction codes.^3,5,6,9^

**Fig. 5:**
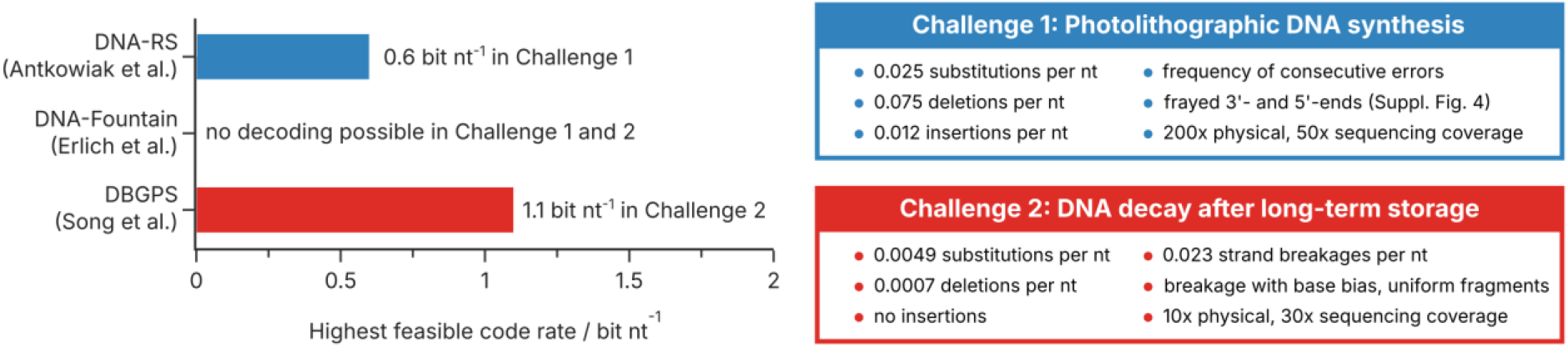
Challenge definitions and the performance of representative error-correction codes. Highest feasible code rates (i.e., code rate at which the 51.6 kB file was perfectly decoded in all three runs, see Methods) achieved in the two challenges by DNA-RS (Antkowiak et al.)^12^, DNA-Fountain (Erlich et al.)^3^, and DBGPS (Song et al.)^19^. Note that the benchmarking did not include full optimization of all (hyper-)parameters of the codecs (see Methods). The error rates, coverages, and specifics of the challenges on photolithographic synthesis (blue) and DNA decay (red) are defined on the right, and also specified in the Methods section.

Taken together, these benchmarking results underline the necessity of error-correction codes to adapt to the challenges posed by the new additions to the state-of-the-art workflow. This is highlighted by the stark performance differences between the tested codecs, despite the use of mostly default (hyper-)parameters. Moreover, these benchmarking results also outline the current state-of-the-art in error-correction coding for photolithographic synthesis and DNA decay, providing means to compare codec performance.

## Discussion

This study characterized the errors and biases present in two new additions to the state-of-the-art workflow in DNA data storage – photolithographic synthesis and DNA decay – and highlighted the associated challenges for error-correction codes. Photolithographic synthesis challenges error-correction codes due the highly erroneous DNA it produces (e.g., > 0.1 errors per nt), necessitating sufficient within-sequence redundancy as well as optimal use of the available redundancy in the sequencing data (e.g., by clustering). For DNA decay, the challenge is two-fold: while error-correction codes must either first reassemble full-length sequences or directly use partial sequences for decoding, our analysis revealed a strong bias stemming from the sequencing workflow used for recovering the oligo fragments after decay. For the latter, we identified 3’-end repair with 3’-phosphodiesterases such as APE1, combined with a single-stranded sequencing preparation workflow, as optimal for ensuring uniform, unbiased recovery of oligo fragments. This is comparable to the state-of-the-art for ancient DNA analysis, in which the use of single-stranded protocols and endonucleases for end repair has also led to improved recovery.^41–43^ To this end, transferring these protocols for ancient DNA recovery to DNA data storage could yield further improvements beyond the results by Meiser et al.^17^, for example by optimizing size selection.^43^

Importantly, our results on the errors and biases in photolithographic synthesis and DNA decay also provide a thorough reference for developers of suitable error-correction codes. For example, our results on DNA decay confirm fundamental assumptions for the DNA-based torn paper channel^30^ (e.g., uniform breakage probability, homogeneous base coverage) and suggest the results by Song et al.^19^ might have been limited by the workflow-induced bias in their sequencing data. Moreover, our benchmarking results for the two challenges presented in this work highlight the current state-of-the-art, and provide standardized, realistic scenarios to measure codec performance. Accordingly, we provide both challenges for use as benchmarks – either online at https://dt4dds.ethz.ch or implemented in C++ at https://github.com/fml-ethz/dt4dds-challenges – and are tracking progress towards these challenges on a leaderboard. Because of the combinatorial complexity of both challenges, we also establish requirements on input size and run time in our challenges to ensure codecs can scale to relevant data sizes (for more information, see https://dt4dds.ethz.ch/challenge).

One of the limitations of this study is the omission of nanopore sequencing as an emerging sequencing technology in DNA data storage. Its implementation presents another challenging addition to the state-of-the-art workflow in DNA data storage, both due to its high error rate and the concatenation of oligos it often requires for efficient operation.^44–46^ In addition, our benchmarking results provide only a lower bound on the performance of DBGPS^19^ and DNA-RS^12^, as no comprehensive optimization of these codecs’ parameters was performed. Nonetheless, we believe this study provides valuable data and tools for tackling current challenges in DNA data storage, especially when considering their combination in the future: enabling data storage with low-cost, photolithographic DNA synthesis and sufficient resilience against DNA decay for long-term storage.

## Methods

### Datasets for photolithographic synthesis

The sequencing data originating from the photolithographic syntheses by Antkowiak et al.^12^ (*File 1, File 2*, and *File 3*) and Lietard et al.^11^ (“2SZ” as *Normal*, “Capped 2SZ” as *Capped*, “4SZ” as *Spaced*, and “CB_120” as *Denser*) were used for analysis. The sequencing data was obtained from the deposited files on figshare^47^ and the European Nucleotide Archive^48^ (PRJEB43002), with the exception of *File 3* (Antkowiak et al.) and *Denser* (Lietard et al.) which the authors did not deposit due to their size and were thus obtained from the corresponding authors directly.

In the study by Antkowiak et al.^12^, three files of different size were synthesized (File 1: 16,383 sequences, File 2: 49,148 sequences, File 3: 196,595 sequences; each with 60 nt). In all cases, the samples were prepared for sequencing with the Accel-NGS 1S Plus DNA Library Kit (Swift Biosciences) and ultimately sequenced on an Illumina NextSeq Sequencer (single-end 75 bp).^12^ In the study by Lietard et al.^11^, the error profiles of four syntheses was analysed. In the first set of three experiments (*Normal, Capped*, and *Spaced*), the same sequences (19,794 sequences with 67 nt) were synthesized while changing the experimental conditions on the synthesizer (i.e., introducing a capping step or increasing the spacing of light features).^11^ In the fourth synthesis (*Denser*), a larger file (382,536 sequences with 76 nt) was synthesized by decreasing the spacing of light features to the minimum. In all four cases, the samples were prepared for sequencing with the Accel-NGS 1S Plus DNA Library Kit (Swift Biosciences) and ultimately sequenced on an Illumina MiSeq Sequencer (paired-end 150 bp).^11^

### Datasets for DNA decay

The sequencing data from Meiser et al.^17^ (“S3” as *unaged*, “S2” as *aged*, and “S1” as *repaired*) and Song et al.^19^ (“P10” as *unaged*, “HT228” as *28 d*, “HT4” as *56 d*, and “HT5” as *70 d*) were used for analysis. All sequencing data was obtained from the deposited files on figshare.^49–51^

In the study by Meiser et al.^17^, an oligo pool synthesized by Twist Bioscience (7,373 sequences with 150 nt, of which 40 nt are primer adapters) was used to test the effect of repair enzymes on the recovery of oligo fragments after aging. Three samples were prepared for sequencing: the oligo pool as-received (*unaged*), after aging for four weeks at 30°C (*aged*), and after aging and subsequent treatment with an enzyme mix (*repaired*).^17^ All samples were prepared for sequencing with the Accel-NGS 1S Plus DNA Library Kit (Swift Biosciences) and ultimately sequenced on an Illumina iSeq 100 Sequencer (paired-end 150 bp).^17^ In the study by Song et al.^19^, an oligo pool synthesized by Twist Bioscience (210,000 sequences with 200 nt, of which 36 nt were primer adapters) was used to validate the performance of their de novo strand assembly for data recovery after aging. Samples from the pool were prepared at four different timepoints during aging at 70°C: as-received (*unaged*), after 28 days (*28 d*), 56 days (*56 d*), and 70 days (*70 d*). All samples were prepared for sequencing with the Illumina TruSeq DNA PCR-Free Library Preparation Kit (Illumina, USA) and sequencing on an Illumina HiSeq Sequencer (paired-end 150 bp).^19^

### Post-processing of sequencing data

The sequencing datasets were filtered to remove reads from sequencing adapters using the script *bbduk*.*sh* from the BBMap suite^52^ (v39.01), using the options “ktrim=r k=23 mink=11 hdist=1 tpe tbo”. The paired reads for the sequencing data on DNA decay were then merged using NGmerge^53^ (v0.3), using the options “-m 10 -d -e 10” to merge overlaps longer than 10 nucleotides with dovetails. Merging of sequencing reads was required due to the length of the design sequences exceeding the read length (150 nt). To map the sequencing data to the reference sequences, *bbmap*.*sh* from the BBMap suite^52^ (v39.01) was used to create a sequence alignment map. Based on the tool’s recommendations, the options “vslow k=8 maxindel=200 minratio=0.1” were used to map sequencing reads with high sensitivity. For the sequencing data on DNA decay, the flag “local=t” was added to obtain local alignments.

### Analysis of sequencing data

To analyse the errors and breakage patterns in the sequencing data, the obtained sequence alignment maps were parsed with the analysis scripts implemented in the Python package DT4DDS by Gimpel et al.^9^ (v1.1). In short, sequence alignments were filtered based on alignment quality (85% similarity for decay datasets, 70% similarity for photolithography datasets), and up to two million reads were separated into reads originating from the sense or antisense directions of the design sequences. In both cases, 3’-overhangs consisting of C and T – originating from the Accel-NGS 1S Plus DNA Library Kit (Swift Biosciences)^26^ employed in the datasets by Meiser et al.^17^ and Antkowiak et al.^12^ – were removed from the alignment. For each read, the occurrence of substitution, insertion and deletion errors, the length of the aligned sequence, and the positions and nucleobases of 3’- and 5’-breaks, among others, were recorded. These statistics were aggregated from all analysed reads from a sequencing dataset and then used to generate the data presented in this work. Full documentation is provided with the GitHub repository and the Python package DT4DDS.^9,54^

### Implementation of Challenge 1: Photolithographic DNA Synthesis

Based on the analysis of error rates and biases in the sequencing data for photolithographic synthesis (see above), a realistic scenario involving the use of photolithographically synthesized DNA for DNA-of-things^13^ was conceptualized. The scenario assumes error rates of 0.075 nt^-1^ for deletions, 0.012 nt^-1^ for insertions, and 0.025 nt^-1^ for substitutions, leading to distributions of consecutive errors and errors per read as shown in Fig. 2d+e. The sequence coverage is biased, assuming a lognormal distribution with a mean of 1 and a standard deviation of 0.44 (see Fig. 2b), and the sequence length is restricted to 80 nucleotides. In addition, the beginning and end of each sequence are truncated according to the observed fragment distribution (see Supplementary Fig. 4). During sampling, a mean physical coverage of 200 oligos per design sequence is assumed, from which a mean sequencing depth of 50 reads per design sequence is derived. The accuracy of the challenge in reproducing the error patterns from photolithographic synthesis is highlighted in Supplementary Fig. 6. The challenge is implemented in C++ (https://github.com/fml-ethz/dt4dds-challenges) as well as available online at https://dt4dds.ethz.ch.

### Implementation of Challenge 2: DNA Decay after long-term storage

Based on the analysis of error rates and biases in the sequencing data for DNA decay (see above), a realistic scenario involving the use of commercially synthesized DNA for long-term, high-density archival storage was conceptualized. The scenario assumes error rates of high-fidelity DNA synthesis and amplification, leading to mean error rates of 0.0007 nt^-1^ for deletions, and 0.0049 nt^-1^ for substitutions. The sequence coverage is biased, assuming a lognormal distribution with a mean of 1 and a standard deviation of 0.30 after synthesis. For storage, a mean initial physical coverage of only 10 oligos per design sequence is assumed. During storage, strand cleavage occurs uniformly along the oligos at a rate of 0.023 breaks nt^-1^, with a base bias towards G (48.8%) and A (39.0%) over C (4.9%) and T (7.3%). Assuming an optimized recovery workflow based on Meiser et al.^17^ with 3’-end repair, all fragments are recovered, with a bias towards fragments longer than 50 nt due to the size cut-off of the purification step (see Supplementary Fig. 3). In addition, all fragments are 3’-tailed with a variable number of C and T nucleotides, to simulate the workflow using the Accel-NGS 1S Plus DNA Library Kit (Swift Biosciences)^26^. These fragments are then sequenced with a mean sequencing depth of 30 reads per design sequence. The accuracy of the challenge in reproducing the error patterns from DNA decay is highlighted in Supplementary Fig. 7.

The challenge is implemented in C++ (https://github.com/fml-ethz/dt4dds-challenges) as well as available online at https://dt4dds.ethz.ch.

### Benchmarking codecs for challenges

Using the implemented challenges for photolithographic synthesis and DNA decay (see above), three representative codecs for DNA data storage – DNA-Fountain by Erlich et al.^3,55^, DBGPS with fountain codes by Song et al.^19,40,40^, and DNA-RS with Reed-Solomon codes by R. Heckel^12,16,38^ – were benchmarked. A compressed SEM micrograph of DNA encapsulated in silica nanoparticles (see Supplementary Fig. 5, file size 51,632 bytes) was used as input file for all tests. For all codecs, default (hyper-)parameters were used, with the exception of sequence count and sequence length. Importantly, no optimization of each codec’s (hyper-)parameters was performed. The number of sequences was modified to obtain various code rates in increments of 0.1 bit nt^-1^. In addition, the length of sequences was adjusted from its default value for the photolithography challenge to conform to the constraint on sequence length during synthesis. For a complete overview of the parameters used for each codec, see Supplementary Tables 1-3.

The determination of the codecs’ code rates neglects constant regions (e.g., constant adapters added to aid strand reassembly or PCR primers) to simplify comparison. For each code rate, the input file was encoded into DNA sequences once, and these design sequences were subsequently used for running the challenges. Each combination of challenge and code rate was run, and decoding attempted, three times to ensure consistent results. The highest code rate for which all three runs led to successful recovery of the input file (i.e., byte-by-byte identity of the original and recovered file) was reported as the codec’s achievable code rate. Full documentation is provided with the GitHub repository.

## Acknowledgments

This project was financed by the European Union’s Horizon 2020 Program, FET-Open: DNA-FAIRYLIGHTS, grant agreement no. 964995, and the European Union’s Horizon EIC Pathfinder Challenge Program: DiDAX, Grant Agreement No. 101115134. Data analysis and simulations were performed on the Euler cluster operated by the High-Performance Computing group at ETH Zürich. Figures were partially created with BioRender.com.

## Author contributions

R.N.G. and R.H. initiated and supervised the project with input from W.J.S. A.L.G. developed the code, performed data analysis, prepared illustrations, and wrote the manuscript with input and approval from all authors.

## Competing interests

The authors declare no competing financial interest.

## Data availability

The sequencing data used in this study is available from Antkowiak et al.^12^ (via figshare^47^), Lietard et al.^11^ (via ENA, accession code PRJEB43002), Meiser et al.^17^ (via figshare^49^), and Song et al.^19^ (via figshare^50,51^). No new sequencing data was generated for this study. The data files derived from the sequencing data and used for the analysis are provided with the code (see code availability statement).

## Code availability

The code and data for error analysis is deposited in the public GitHub repository at https://github.com/fml-ethz/dt4dds-challenges_notebooks. The implementation of the challenges used for benchmarking are deposited in in the public GitHub repository at https://github.com/fml-ethz/dt4dds-challenges.

## Additional Information

Supplementary Information is available for this paper.

## Notes

https://doi.org/10.6084/m9.figshare.c.5128901.v1

https://doi.org/10.6084/m9.figshare.21070684.v1

https://doi.org/10.6084/m9.figshare.17193170.v2

https://doi.org/10.6084/m9.figshare.17192639.v1

https://www.ebi.ac.uk/ena/browser/view/PRJEB43002

https://dt4dds.ethz.ch/challenge

### Competing Interest Statement

The authors have declared no competing interest.

## References

1. Ceze, L., Nivala, J. & Strauss, K. Molecular digital data storage using DNA. Nat. Rev. Genet. 2019 208 20, 456–466 (2019).

2. Grass, R. N., Heckel, R., Puddu, M., Paunescu, D. & Stark, W. J. Robust Chemical Preservation of Digital Information on DNA in Silica with Error-Correcting Codes. Angew. Chem. Int. Ed. 54, 2552–2555 (2015).

3. Erlich, Y. & Zielinski, D. DNA Fountain enables a robust and efficient storage architecture. Science 355, 950–954 (2017).

4. Goldman, N. et al. Towards practical, high-capacity, low-maintenance information storage in synthesized DNA. Nat. 2013 4947435 494, 77–80 (2013).

5. Ping, Z. et al. Towards practical and robust DNA-based data archiving using the yin–yang codec system. Nat. Comput. Sci. 2022 24 2, 234–242 (2022).

6. Welzel, M. et al. DNA-Aeon provides flexible arithmetic coding for constraint adherence and error correction in DNA storage. Nat. Commun. 14, 628 (2023).

7. Church, G. M., Gao, Y. & Kosuri, S. Next-generation digital information storage in DNA. Science 337, 1628 (2012).

8. Yu, M. et al. High-throughput DNA synthesis for data storage. Chem. Soc. Rev. 53, 4463–4489 (2024).

9. Gimpel, A. L., Stark, W. J., Heckel, R. & Grass, R. N. A digital twin for DNA data storage based on comprehensive quantification of errors and biases. Nat. Commun. 14, 6026 (2023).

10. Matange, K., Tuck, J. M. & Keung, A. J. DNA stability: a central design consideration for DNA data storage systems. Nat. Commun. 12, 1358 (2021).

11. Lietard, J. et al. Chemical and photochemical error rates in light-directed synthesis of complex DNA libraries. Nucleic Acids Res. 49, 6687–6701 (2021).

12. Antkowiak, P. L. et al. Low cost DNA data storage using photolithographic synthesis and advanced information reconstruction and error correction. Nat. Commun. 11, 5345 (2020).

13. Koch, J. et al. A DNA-of-things storage architecture to create materials with embedded memory. Nat. Biotechnol. 2019 381 38, 39–43 (2019).

14. Meiser, L. C. et al. Synthetic DNA applications in information technology. Nat. Commun. 13, 352 (2022).

15. Lim, C. K., Nirantar, S., Yew, W. S. & Poh, C. L. Novel Modalities in DNA Data Storage. Trends Biotechnol. 39, 990–1003 (2021).

16. Meiser, L. C. et al. Reading and writing digital data in DNA. Nat. Protoc. 2019 151 15, 86–101 (2019).

17. Meiser, L. C. et al. Information decay and enzymatic information recovery for DNA data storage. Commun. Biol. 5, 1–9 (2022).

18. Heckel, R., Mikutis, G. & Grass, R. N. A Characterization of the DNA Data Storage Channel. Sci. Rep. 2019 91 9, 1–12 (2019).

19. Song, L. et al. Robust data storage in DNA by de Bruijn graph-based de novo strand assembly. Nat. Commun. 2022 131 13, 1–9 (2022).

20. Organick, L. et al. Probing the physical limits of reliable DNA data retrieval. Nat. Commun. 2020 111 11, 1–7 (2020).

21. Filges, S., Mouhanna, P. & Ståhlberg, A. Digital Quantification of Chemical Oligonucleotide Synthesis Errors. Clin. Chem. 67, 1384–1394 (2021).

22. Xu, C. et al. Electrochemical DNA synthesis and sequencing on a single electrode with scalability for integrated data storage. Sci. Adv. 7, eabk0100 (2021).

23. Nguyen, B. H. et al. Scaling DNA data storage with nanoscale electrode wells. Sci. Adv. 7, eabi6714 (2021).

24. Chen, Y.-J. et al. Quantifying molecular bias in DNA data storage. Nat. Commun. 2020 111 11, 1–9 (2020).

25. Gao, Y., Chen, X., Qiao, H., Ke, Y. & Qi, H. Low-Bias Manipulation of DNA Oligo Pool for Robust Data Storage. ACS Synth. Biol. 9, 3344–3352 (2020).

26. Swift Biosciences. ACCEL-NGS® 1S Plus DNA Library Kit, Protocol for Cat. Nos. 10024 and 10096 (2018).

27. Illumina Inc. TruSeq DNA PCR-Free Reference Guide, Document #1000000039279 (2017).

28. Beckman Coulter. AMPure XP: Manual or Automated Purification and Clean-up, Document #AAG-4464DS12.18 (2019).

29. Mikutis, G., Schmid, L., Stark, W. J. & Grass, R. N. Length-dependent DNA degradation kinetic model: Decay compensation in DNA tracer concentration measurements. AIChE J. 65, 40–48 (2019).

30. Bar-Lev, D., Marcovich, S., Yaakobi, E. & Yehezkeally, Y. Adversarial Torn-Paper Codes. IEEE Trans. Inf. Theory 69, 6414–6427 (2023).

31. Bruce, A. et al. Molecular Biology of the Cell: Seventh International Student Edition with Registration Card. (W.W. Norton & Company, 2022).

32. Hegde, M. L., Hazra, T. K. & Mitra, S. Early steps in the DNA base excision/single-strand interruption repair pathway in mammalian cells. Cell Res. 18, 27–47 (2008).

33. Krokan, H. E. & Bjørås, M. Base Excision Repair. Cold Spring Harb. Perspect. Biol. 5, a012583 (2013).

34. Suh, D., Wilson, D. M., III & Povirk, L. F. 3′-Phosphodiesterase activity of human apurinic/apyrimidinic endonuclease at DNA double-strand break ends. Nucleic Acids Res. 25, 2495–2500 (1997).

35. Lindahl, T. Instability and decay of the primary structure of DNA. Nat. 1993 3626422 362, 709–715 (1993).

36. Mitchell, D., Willerslev, E. & Hansen, A. Damage and repair of ancient DNA. Mutat. Res. Mol. Mech. Mutagen. 571, 265–276 (2005).

37. Shapiro, R. Damage to DNA Caused by Hydrolysis. in Chromosome Damage and Repair (eds. Seeberg, E. & Kleppe, K.) 3–18 (Springer US, New York, NY, 1981).

38. Heckel, R. reinhardh/dna_rs_coding: Error correction scheme for storing information on DNA using Reed Solomon codes. GitHub (2021).

39. Darestani, M. Z. & Heckel, R. MLI-lab/noisy_dna_data_storage: Data recovery from millions of noisy reads. Zenodo 10.5281/zenodo.4044459 (2020).

40. Song, L. DBGPS (Python) and fountain codes for robust data storage in DNA. Zenodo 10.5281/zenodo.6833784 (2022).

41. Briggs, A. W. et al. Removal of deaminated cytosines and detection of in vivo methylation in ancient DNA. Nucleic Acids Res. 38, e87 (2010).

42. Orlando, L. et al. Ancient DNA analysis. Nat. Rev. Methods Primer 1, 1–26 (2021).

43. Gansauge, M.-T., Aximu-Petri, A., Nagel, S. & Meyer, M. Manual and automated preparation of single-stranded DNA libraries for the sequencing of DNA from ancient biological remains and other sources of highly degraded DNA. Nat. Protoc. 15, 2279–2300 (2020).

44. Organick, L. et al. Random access in large-scale DNA data storage. Nat. Biotechnol. 2018 363 36, 242–248 (2018).

45. Delahaye, C. & Nicolas, J. Sequencing DNA with nanopores: Troubles and biases. PLOS ONE 16, e0257521 (2021).

46. Lopez, R. et al. DNA assembly for nanopore data storage readout. Nat. Commun. 2019 101 10, 1–9 (2019).

47. Antkowiak, P. Low Cost DNA Data Storage Using Photolithographic Synthesis and Advanced Information Reconstruction and Error Correction. figshare 10.6084/m9.figshare.c.5128901.v1 (2020).

48. Lietard, J. DNA_photolithography_oligo, Project PRJEB43002. European Nucleotide Archive.

49. Meiser. Sequencing data of Meiser et al. Comms. Biol. 2022. figshare 10.6084/m9.figshare.21070684.v1 (2022).

50. Song, L. Accelerated aging samples of 70-°C for 0 and 28 days. figshare 10.6084/m9.figshare.17193170.v2 (2021).

51. Song, L. Accelerated aging samples of 70-°C for 56 and 70 days. figshare 10.6084/m9.figshare.17192639.v1 (2021).

52. Bushnell, B. BBMap: A Fast, Accurate, Splice-Aware Aligner. https://www.osti.gov/biblio/1241166 (2014).

53. Gaspar, J. M. NGmerge: merging paired-end reads via novel empirically-derived models of sequencing errors. BMC Bioinformatics 19, 536 (2018).

54. Gimpel, A. fml-ethz/dt4dds: v1.0.0. Zenodo 10.5281/zenodo.8329037 (2023).

55. Erlich, Y. TeamErlich/dna-fountain. GitHub (2024).

