## Supplementary Figures and Tables for "Challenges for error-correction coding in DNA data storage: photolithographic synthesis and DNA decay"

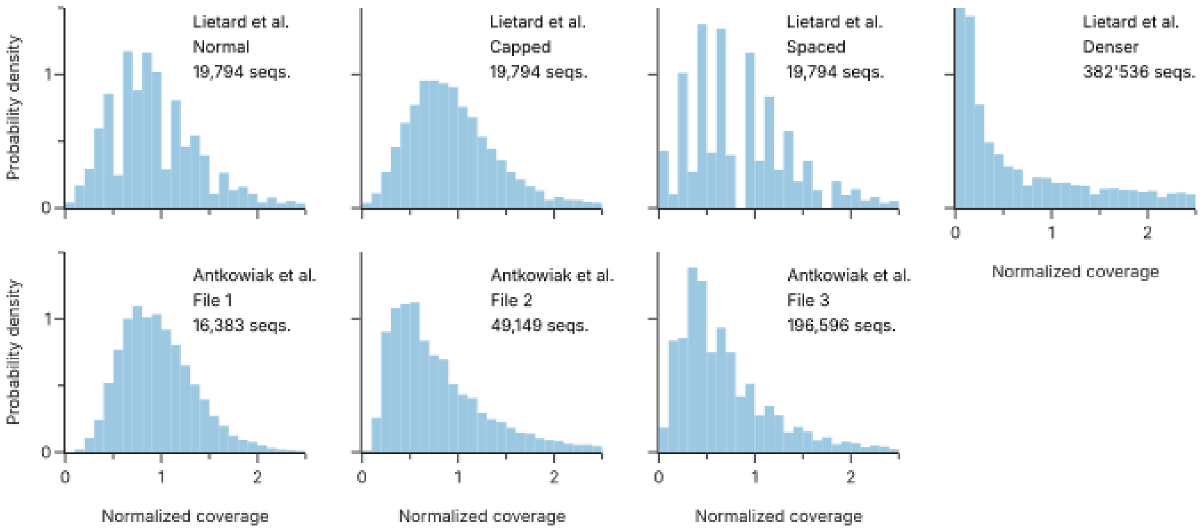


**Supplementary Fig. 1: Normalized sequences coverage for photolithographic syntheses.** The homogeneity of the oligo pools synthesized by Lietard et al.[1] (top row) and Antkowiak et al.[2] (bottom row) shows an increase in skewness and bias with the number of sequences synthesized in parallel. Due to the low sequencing depth, especially in the datasets by Lietard et al.[1], the data is non-continuous and therefore the histograms appear non-uniform.

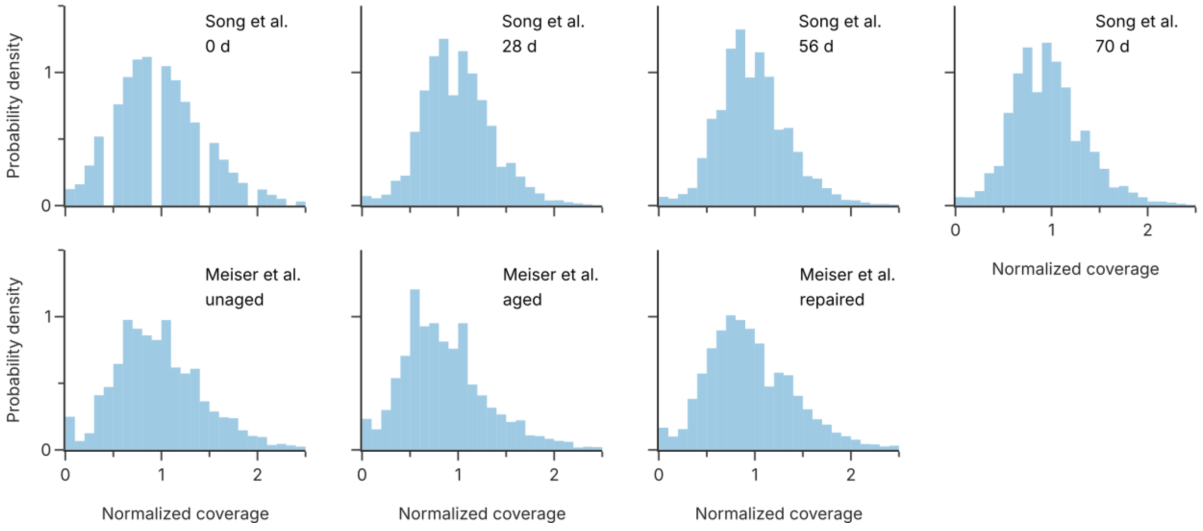


**Supplementary Fig. 2: Normalized sequences coverage for DNA decay.** The homogeneity of the oligo pools synthesized by Song et al.[3] (top row) and Meiser et al.[4] (bottom row) shows no increase in skewness and bias with aging.

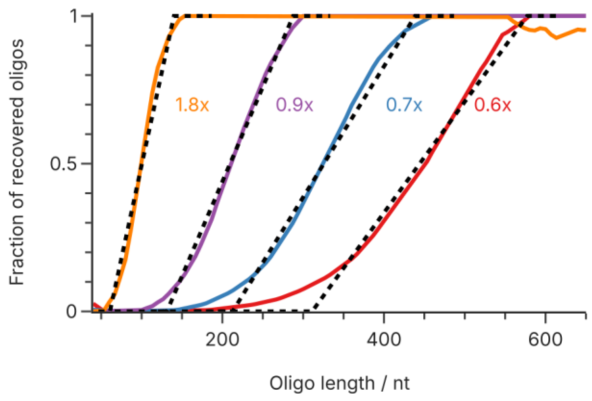


**Supplementary Fig. 3: Recovery rate during bead-based clean-up as a function of oligo length and bead ratio.** The oligo recovery workflows by Song et al.[3] and Meiser et al.[4] use bead-based clean-up to remove excess adapters after ligation. At different bead ratios (1.8x to 0.6x), the proportion of oligos recovered depends strongly on their individual lengths. This relationship is reasonably well approximated by a piecewise linear function (dotted lines) with a lower cut-off and upper threshold length. The raw data is based on the experimental data provided by Beckman Coulter for AMPure XP beads.[5]

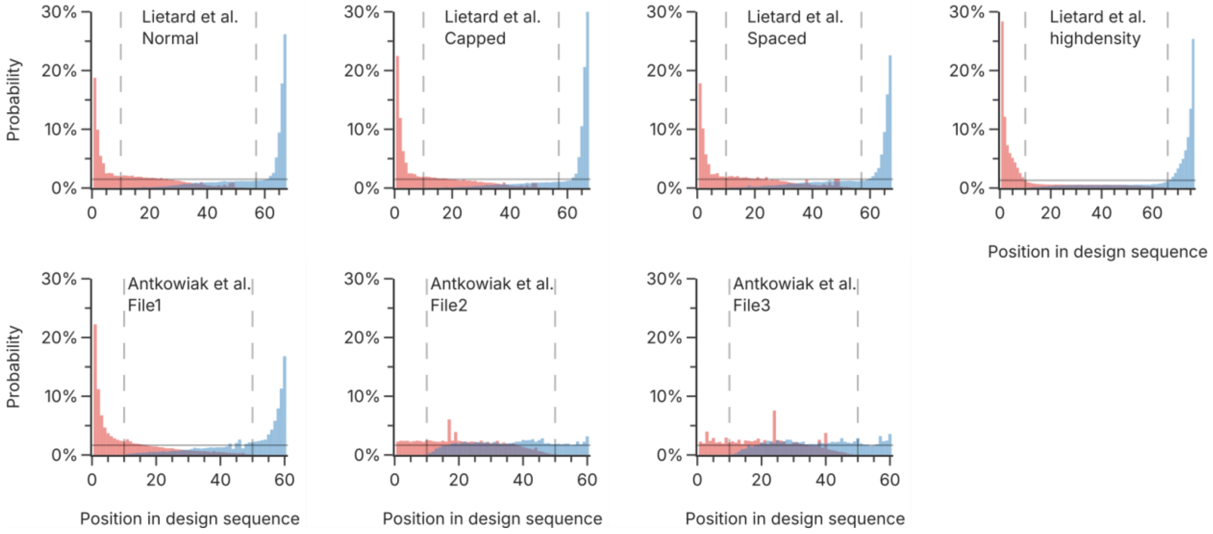


**Supplementary Fig. 4: Positional distribution of 3'- (blue) and 5'-ends (red) for photolithographic syntheses.** The fragments in the sequencing data by Lietard et al.[1] (top row) and Antkowiak et al.[2] (bottom row) show a clear trend towards truncation at both the 3'- and 5'-ends, as long as 10 nt at each end. The datasets of File 2 and File 3 of Antkowiak et al.[2] do not follow this trend, as the positional distribution is even along the design sequence. Dotted lines indicate the filtering of short sequences by the analysis pipeline (i.e., reads shorter than 10 nt are removed). The solid horizontal line indicates an equal probability along the design sequence.

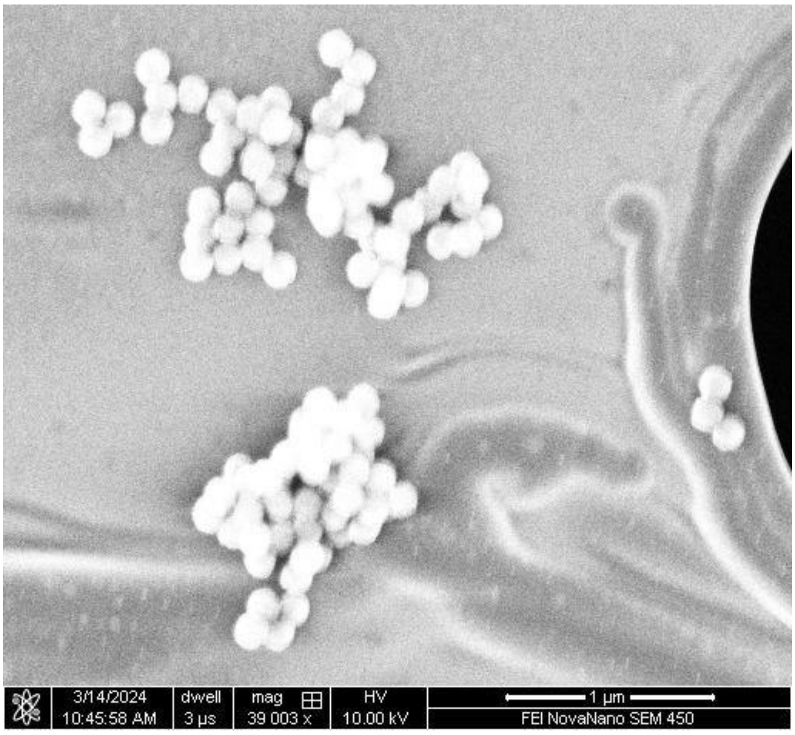


**Supplementary Fig. 5: SEM micrograph of silica nanoparticles encapsulating DNA.** This image, compressed into a zip file of 51,632 bytes, was used as input for the codecs benchmarked for the two challenges described in the main text.

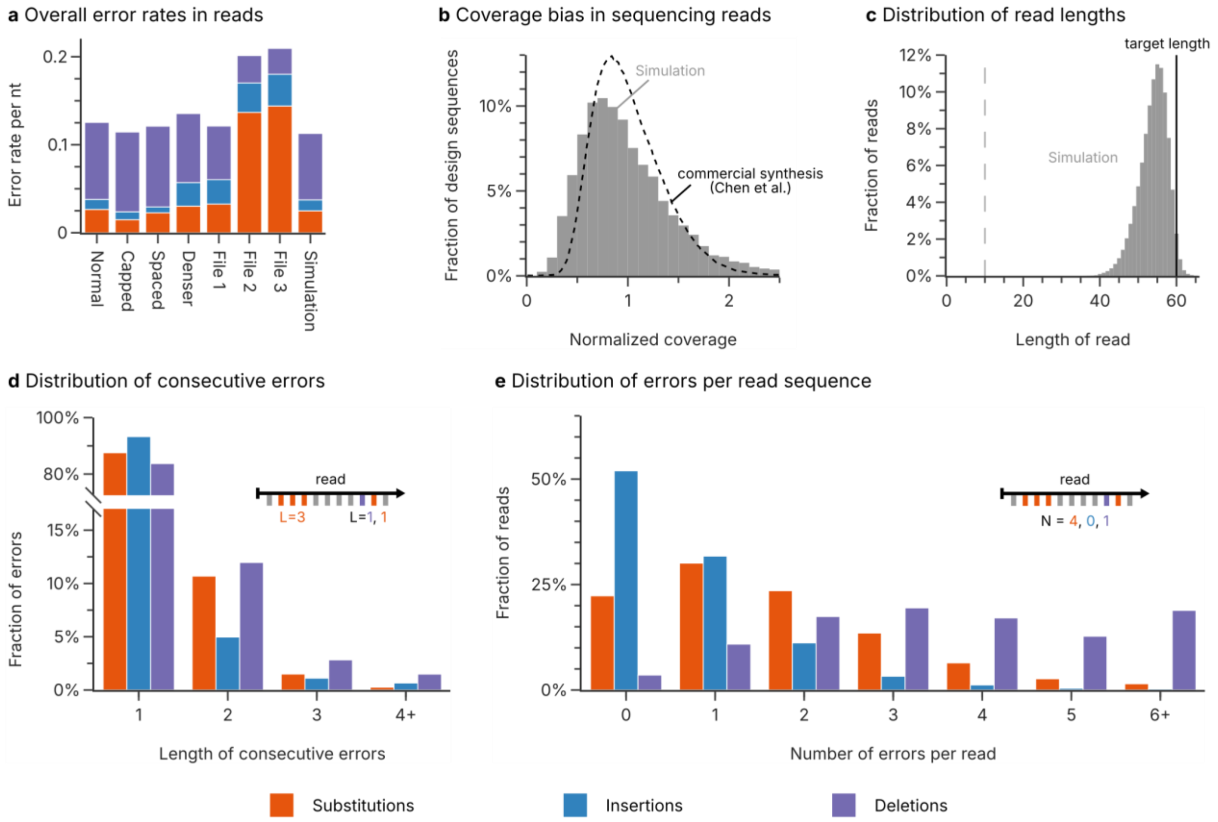


**Supplementary Fig. 6: Analysis of photolithographic synthesis as simulated by Challenge 1. (a)** Overall rate of substitution (orange), insertion (blue), and deletion (purple) errors in the sequencing datasets for photolithographic syntheses by Lietard et al.[1], Antkowiak et al.[2] and the simulation. The error rates represent the median error rate across the length of the sequence, to minimize the effect of the low-diversity regions at the start and end of the sequences. **(b)** Distribution of the sequence coverage in the simulated dataset compared to the coverage distribution for a commercial synthesis by material deposition (Twist Biosciences, data by Chen et al.[6]). **(c)** Length distribution of the reads in the simulated dataset compared to the length of the design sequences (solid line). Only the segment of the read which aligned to the reference sequence is considered. Reads smaller than 10 nucleotides were discarded during mapping of the sequencing data (dotted line). **(d)** The length of consecutive of substitution (orange), insertion (blue), and deletion (purple) errors in the simulated sequencing dataset. **(e)** The number of substitution (orange), insertion (blue), and deletion (purple) errors in the sequencing reads in the simulated dataset. The data in this figure can be compared to Fig. 2 in the main text to validate the accuracy of the digital model of photolithographic synthesis.

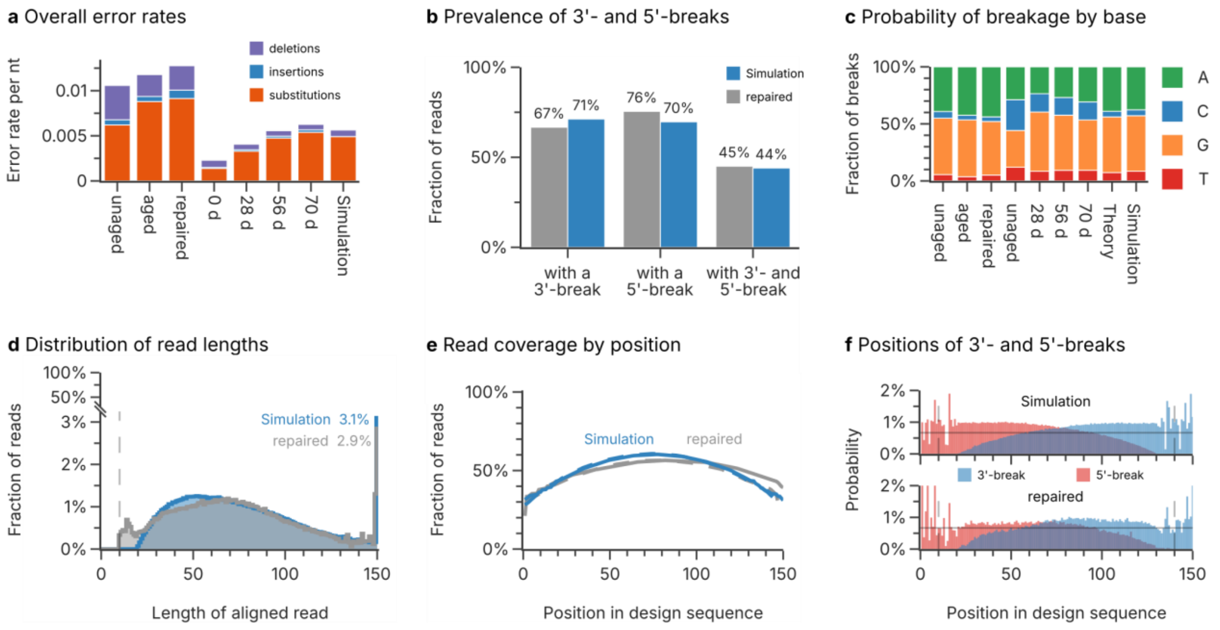


**Supplementary Fig. 7: Analysis of DNA decay during aging as simulated by Challenge 2. (a)** Overall rate of substitution (orange), insertion (blue), and deletion (purple) errors in the sequencing datasets by Meiser et al.[4], Song et al.[3], and the simulation. **(b)** Fraction of reads with different break patterns in the repaired dataset by Meiser et al.[4] (gray) and the simulated dataset (blue). **(c)** Distribution of nucleobases (green: A, blue: C, orange: G, red: T) involved in strand cleavage in the sequencing datasets by Meiser et al.[4], Song et al.[3], the simulation, as well as the theoretical distribution expected from reaction rates. Experimental distributions are estimated from the type of nucleobase preceding a 5'-break in the sequencing data. **(d)** Histograms of the length distribution of aligned sequencing reads in the repaired dataset by Meiser et al.[4] (gray) and the simulated dataset (blue). The fraction of reads with full length in each dataset are given as a percentage. **(e)** Read coverage by position in the design sequence of the repaired dataset by Meiser et al.[4] (gray) and the simulation (blue). The reads are separated by their direction into sense (solid lines) and antisense (dashed lines) directions. A value of 100% corresponds to every read containing a specific position. **(f)** Positional distributions of 3'- (blue) and 5'-breaks (red) in the repaired dataset by Meiser et al.[4] (bottom) and the simulated dataset(top). The horizontal solid line denotes the expected breakage probability if decay occurs uniformly. The data in this figure can be compared to Figs. 3+4 in the main text to validate the accuracy of the digital model of DNA decay during long-term storage.

#### Supplementary Tables

**Supplementary Table 1: Settings for the DNA-RS codec by Antkowiak et al.[2] used in benchmarking.**

| Challenge | Code rate | Length | Redundancy | # Sequences | Success |
|---|---|---|---|---|---|
| Photolithographic Synthesis | 0.80 bit $nt^{-1}$ | 72 | 6 | 7200 | 0/3 |
| | 0.70 bit $nt^{-1}$ | 72 | 6 | 8200 | 0/3 |
| | 0.60 bit $nt^{-1}$ | 72 | 6 | 9500 | 3/3 |
| | 0.50 bit $nt^{-1}$ | 72 | 6 | 11500 | 3/3 |

**Supplementary Table 2: Settings for the DBGPS codec by Song et al.[3] used in benchmarking.**

| Challenge | Code rate | Chunk size | # Droplets | Success |
|---|---|---|---|---|
| DNA Decay | 1.40 bit $nt^{-1}$ | 35 | 1800 | 0/3 |
| | 1.30 bit $nt^{-1}$ | 35 | 1930 | 0/3 |
| | 1.20 bit $nt^{-1}$ | 35 | 2100 | 1/3 |
| | 1.10 bit $nt^{-1}$ | 35 | 2300 | 3/3 |

**Supplementary Table 3: Settings for the DNA-Fountain codec by Erlich et al.[7] used in benchmarking.**

| Challenge | Code rate | Chunk size | Redundancy factor | Success |
|---|---|---|---|---|
| Photolithographic synthesis | 0.50 bit $nt^{-1}$ | 14 | 1.8 | 0/3 |
| | 0.40 bit $nt^{-1}$ | 14 | 2.5 | 0/3 |
| | 0.30 bit $nt^{-1}$ | 14 | 3.6 | 0/3 |
| | 0.20 bit $nt^{-1}$ | 14 | 5.9 | 0/3 |
| DNA Decay | 0.50 bit $nt^{-1}$ | 32 | 2.4 | 0/3 |
| | 0.40 bit $nt^{-1}$ | 32 | 3.2 | 0/3 |
| | 0.30 bit $nt^{-1}$ | 32 | 4.6 | 0/3 |
| | 0.20 bit $nt^{-1}$ | 32 | 7.5 | 0/3 |

### References


1. Lietard, J. *et al.* Chemical and photochemical error rates in light-directed synthesis of complex DNA libraries. *Nucleic Acids Res.* **49**, 6687–6701 (2021).
2. Antkowiak, P. L. *et al.* Low cost DNA data storage using photolithographic synthesis and advanced information reconstruction and error correction. *Nat. Commun.* **11**, 5345 (2020).
3. Song, L. *et al.* Robust data storage in DNA by de Bruijn graph-based de novo strand assembly. *Nat. Commun. 2022 131* **13**, 1–9 (2022).
4. Meiser, L. C. *et al.* Information decay and enzymatic information recovery for DNA data storage. *Commun. Biol.* **5**, 1–9 (2022).
5. Beckman Coulter. AMPure XP: Manual or Automated Purification and Clean-up, Document #AAG-4464DS12.18 (2019).
6. Chen, Y.-J. *et al.* Quantifying molecular bias in DNA data storage. *Nat. Commun. 2020 111* **11**, 1–9 (2020).
7. Erlich, Y. & Zielinski, D. DNA Fountain enables a robust and efficient storage architecture. *Science* **355**, 950–954 (2017).
